# KIF5A regulates axonal repair and time-dependent axonal transport of SFPQ granules and mitochondria in human motor neurons

**DOI:** 10.1101/2024.09.06.611684

**Authors:** Irune Guerra San Juan, Jessie Brunner, Kevin Eggan, Ruud F. Toonen, Matthijs Verhage

## Abstract

Mutations in the microtubule binding motor protein, kinesin family member 5A (KIF5A), cause the fatal motor neuron disease, Amyotrophic Lateral Sclerosis. While KIF5 family members transport a variety of cargos along axons, it is still unclear which cargos are affected by *KIF5A* mutations. We generated *KIF5A* null mutant human motor neurons to investigate the impact of KIF5A loss on the transport of various cargoes and its effect on motor neuron function at two different timepoints *in vitro*. The absence of KIF5A resulted in reduced neurite complexity in young motor neurons (DIV14) and significant defects in axonal regeneration capacity at all developmental stages. KIF5A loss did not affect neurofilament transport but resulted in decreased mitochondria motility and anterograde speed at DIV42. More prominently, KIF5A depletion strongly reduced anterograde transport of SFPQ-associated RNA granules in DIV42 motor neuron axons. We conclude that KIF5A most prominently functions in human motor neurons to promote axonal regrowth after injury as well as to anterogradely transport mitochondria and, to a larger extent, SFPQ-associated RNA granules in a time-dependent manner.

## Introduction

Over the last two decades, growing evidence from genetic studies has positioned disruptions to the axonal cytoskeleton as an emerging biological pathway in the pathogenesis of the motor neuron disease, Amyotrophic Lateral Sclerosis (ALS)^1–5^. Motor neurons, and more specifically, lower motor neurons in the spinal cord, rely heavily on their long axonal projections for proper communication between the cell body and the synaptic terminal at the neuromuscular junction^6–8^. The proper maintenance of the axonal structure depends on a variety of components ranging from the axonal initial segment, through microtubule (MT) tracks and actin networks, to MT-based motor proteins for transport^9–12^. Indeed, mutations in axonal transport-related genes have been implicated in various neurological disorders, highlighting the crucial role of long-range intracellular transport mechanisms in maintaining neuronal function. Despite the outward phenotypes resulting from mutations in genes encoding for transport machinery being quite heterogenous, most of these disease-causing mutations are responsible for only a small subset of neurodevelopmental and neurodegenerative diseases. Notably, out of the large superfamily of motor proteins, only four (DCTN1, KIF1A, KIF1C and KIF5A) give rise to neurodegenerative disorders^13^. In recent years, Whole-Genome Sequencing (WGS) approaches have reported autosomal dominant mutations in the *kinesin family member 5A* (*KIF5A*) gene, a member of the kinesin superfamily of motor proteins, to be associated with ALS ^14–19^. Prior to these discoveries and further supporting the relevance of this Kinesin-1 paralog in motor neuron function, mutations in *KIF5A* have also been described to be the cause of other motor neuron diseases, such as Spastic Paraplegia type 10 (SPG10) and Charcot-Marie-Tooth type 2 (CMT2)^13,20–22^. Even though these gene discovery studies have been critical to strengthen the hypothesis that disturbed cytoskeletal dynamics/axonal transport is a major process affected in neurodegenerative diseases, how these genetic variants in KIF5A contribute to compromised motor neuron function requires further examination.

KIF5A, along with KIF5B and KIF5C, is part of the kinesin superfamily of MT-binding motor proteins, Kinesin-1 (KIF5) in mammals^23^.While KIF5A and KIF5C are neuron specific, KIF5B is ubiquitously expressed^24,25^. In rodents, the developmental expression of the KIF5 protein family in the nervous systems appears to be similarly regulated. More specifically, the expression of Kinesin-1 paralogs is the most pronounced at embryonic and early post-natal stages (P1) and is then reduced in adulthood^26,27^. KIF5 motor complexes are tetramers consisting of two kinesin heavy chains (KHC) and two kinesin light chains (KLC). The KHC comprises an N-terminal microtubule-binding and catalytic motor domain, followed by a central stalk region that allows for dimerization and lastly, a C-terminal domain responsible for cargo binding, autoinhibition and MT-sliding^28–30^. Sequentially, regulation of cargo binding to the motor is carried out by the interaction of the C-terminal domain of the KHC with the KLCs and adaptor proteins. Despite the seemingly shared structural and sequence similarity among KIF5 motors, the C-terminal cargo binding domain differs significantly across the three isoforms, which could potentially confer unique functions to each KIF5 in neurons. In fact, KIF5A is the only KIF5 motor found to be mutated in neurodegenerative disorders. ALS-associated *KIF5A* variants all cluster within the C-terminus tail domain of the kinesin, specifically within the 5’ and 3’ splice junctions of Exon 27, which are predicted to result in a novel C-terminal domain. Recent work suggests a possible molecular mechanism whereby the mutant C-terminus domain results in a hyperactive kinesin, which would affect transport of specific cargos that are important for motor neuron survival^31,32^. Cargos known to be trafficked by the KIF5 family members in mammalian neurons are cytoskeletal elements such as neurofilaments, organelles like mitochondria and lysosomes, and granules containing RNA-binding proteins (RBP) among others^33^. However, which of these cargos specifically depend on KIF5A and would therefore be affected by mutant *KIF5A* is still unclear. Identifying these KIF5A-dependent cargos could further our understanding of axonal transport defects in the pathogenesis of ALS and possibly, other motor neuron disorders.

Here, we employed CRISPR/Cas9 gene editing in human motor neurons to study the effect of KIF5A loss on neurite/axonal outgrowth in homeostatic and injury conditions as well as on axonal transport of various cargoes at two distinct time points during development. KIF5A loss resulted in reduced neurite complexity at early stages and severe defects in axonal repair capacity across development. KIF5A loss did not impact axonal neurofilament transport but reduced mitochondrial motility and to a larger extent SFPQ-associated RNA granule transport. Taken together, our data suggest that KIF5A is required for axonal repair, SFPQ-associated granule transport and, to some extent, mitochondria motility in human motor neurons. These findings harbor plausible connections between compromised motor axon function during injury responses and distinct defects in the axonal transport of cargoes such as mitochondria or SFPQ-containing granules.

## Results

### Differential expression of KIF5 paralogs in human neurons and generation of *KIF5A*^null^ human motor neurons

Given that KIF5A is so far the only KIF5 gene in which mutations result in age-associated neurodegenerative disorders, we wondered whether expression of the *KIF5* transcripts varies in the developing and aging human brain. Based on the transcriptomic datasets available from BrainSpan (Allen Brain Atlas), *KIF5A* and *KIF5C* expression are upregulated at postnatal stages across all brain regions analyzed, while *KIF5B* expression is downregulated. (Figure 1a). Similar upregulation of *KIF5A* and downregulation of *KIF5B* was observed in the motor cortex (M1C), while *KIF5C* expression was unchanged over time (Figure 1b).

**Figure 1.**
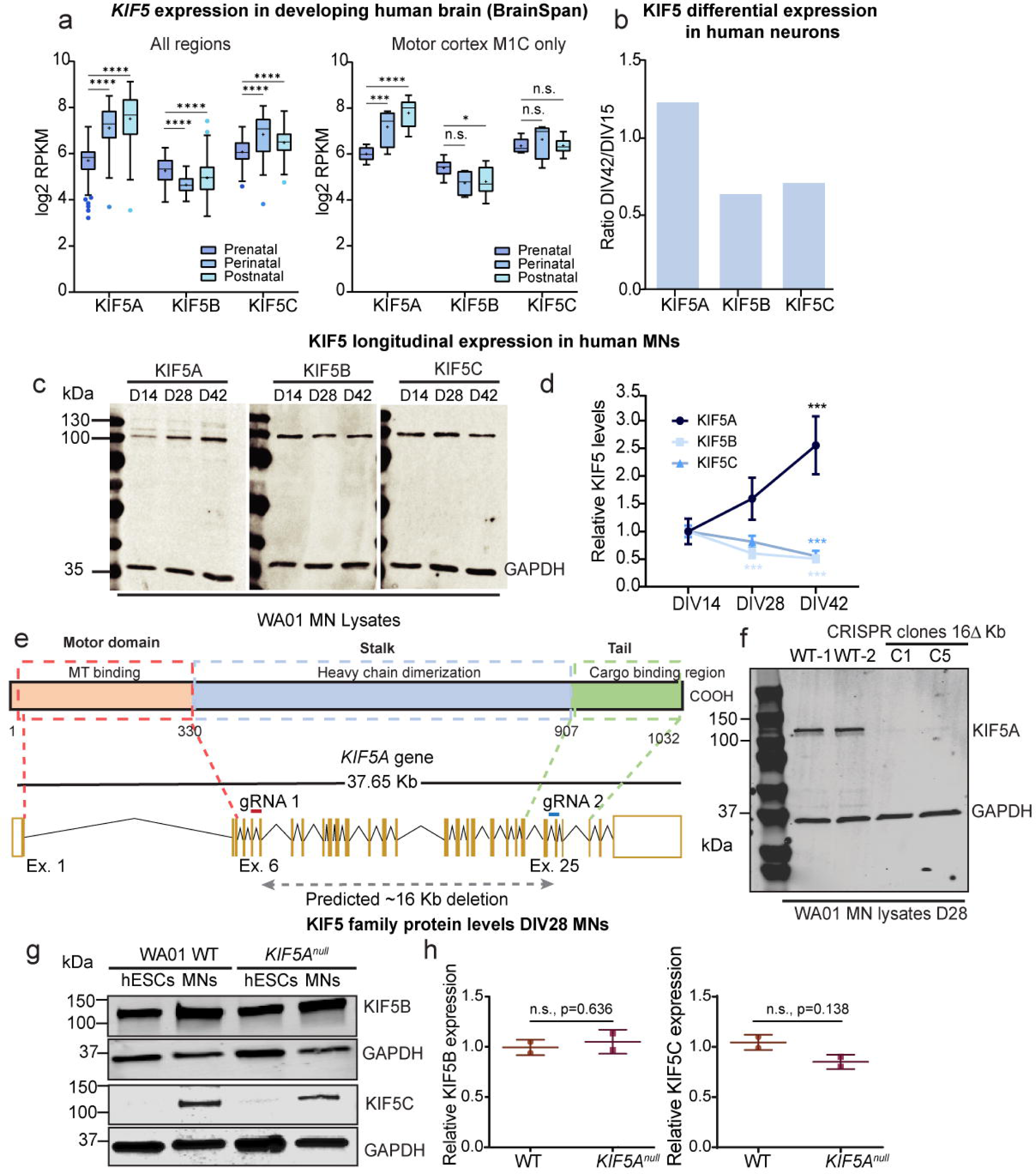
Differential expression of KIF5 paralogs in human neurons and CRISPR/Cas9 editing of *KIF5A*. (A) Relative KIF5 paralog expression in BrainSpan dataset. All brain regions available in BrainSpan are plotted for each KIF5 gene in left panel, and only for motor cortex (M1C) in the right panel. The age groups are divided by prenatal: 1 to 35 post-conception weeks (pcw), perinatal: 37 pcw, 4 months, 10 months and 1 year, and postnatal: 2 to 40 years in both plots. n= 220 donors (all regions) and 9 donors (motor cortex) for the prenatal group, n=75 donors (all regions) and 4 (motor cortex) for perinatal, n=228 (all regions) and n=13 (motor cortex). The 25 pcw age group was excluded in the analysis since it only contained one donor. (B) Differential expression of KIF5 paralogs in proteomic analysis of human neurons at DIV14 and DIV42 (C) Western blot analysis of all three KIF5 paralogs in motor neurons at DIV14, DIV28, and DIV42. (D) Quantification of relative KIF5A paralog expression over time in three independent experiments for KIF5A (E) Schematic representation of the *KIF5A* protein (top) and genomic locus in which gRNAs were targeted to Exon 6 (in red) and Exon 25 (in blue) to generate a large 16 Kb deletion. (F) KIF5A protein levels in induced human motor neurons (MNs) from two selected human embryonic stem cell (hESC, WA01) CRISPR clones including reference protein GAPDH. (G, H) KIF5B and KIF5C protein levels in WT and *KIF5A^null^* motor neurons at Day 28, quantified from Western Blot analysis and normalized to GAPDH (n=2). Data was assessed using a Two-Way ANOVA with Tukey’s multiple comparison test in (A), One-Way ANOVA with Sidak’s multiple comparison test in (D), and a t-test in (H), with significance set to p < 0.05, data are presented in Tukey boxplots in (A) with cross showing mean and central bar median, and in (D,H) as mean with error bars calculated as S.D., with, * p < 0.05, *** p < 0.001, and **** p < 0.0001.

In view of the differential transcriptional regulation of the KIF5 genes *in vivo*, we next studied the expression of the KIF5 proteins in NGN2-induced human neuronal models. iPSCs from three healthy individuals were differentiated into iNeurons using NGN2 expression in combination with dual SMAD and WNT inhibition^34^. iNeurons were harvested 15 (DIV15) and 42 days (DIV42) after induction (Suppl. Figure 1e). Analysis of the protein class “microtubule binding motor protein” showed that many proteins of the Kinesin superfamily were differentially regulated over time in culture (Suppl. Figure 1f). Of note, only the Kinesin-1 paralogs were oppositely regulated: KIF5A was significantly upregulated between DIV15 and 42, while the other two Kinesin-1 members, KIF5B and KIF5C, were downregulated (Figure 1b; Suppl. Figure 1f). A similar pattern was observed in induced lower motor neurons^35^ at DIV14, DIV28 and DIV42 by Western-blotting with KIF5 specific antibodies. Similar to cortical iNeurons, KIF5A levels increased while KIF5B and KIF5C levels decreased between DIV15 and DIV42 (Figure 1c, d).

To investigate the function of *KIF5A* in human motor neurons, we created loss-of-function mutations in the *KIF5A* locus in human embryonic stem cells. We produced a large deletion by targeting Exons 6 and 25 of the *KIF5A* gene that resulted in loss of protein and transcript expression (Figure 1e, f, Suppl. Figure 1a-d). Given the functional redundancy and structural similarity of the Kinesin-1 protein family, protein levels of KIF5B and KIF5C were evaluated for compensatory expression as the result of loss of KIF5A. Western blot analysis at DIV28 showed no significant changes in protein levels of KIF5B and KIF5C in *KIF5A^null^*mutant motor neurons compared to wild-type (WT) controls (Figure 1g,h).

### Early neurite outgrowth deficits in *KIF5A*^null^ human motor neurons

Next, we asked whether the absence of KIF5A affects human motor neuron morphology and synapse density. We performed immunostaining of neurite/cytoskeletal markers in WT and *KIF5A*^null^ motor neurons at DIV14 an DIV42 (Figure 2a). WT neurons showed a gradual increase in MAP2-positive dendrite length and PSD-95-positive synapse number over time in culture (Suppl. Fig. 2a-d). At DIV14, β-III-Tubulin (TUJ1) staining of *KIF5A*^null^ motor neurons showed a significant reduction in total neurite length (Figure 2b-c), accompanied by a decrease in proximal neurite branching as quantified by Sholl analysis in comparison to WT neurons (Figure 2c). In contrast, at DIV42 both total length of MAP2-positive dendrites and SMI32-positive neurites (Figure 2e-f) as well as branching were normal in KIF5A*^null^* mutant motor neurons (Figure 2g). The PSD95 puncta density was similar between *KIF5A*^null^ and WT controls (Suppl. Figure 2b,c). Therefore, KIF5A expression in motor neurons is necessary for normal initial neurite outgrowth and arborization but is dispensable for branching and synapse formation in older motor neurons.

**Figure 2.**
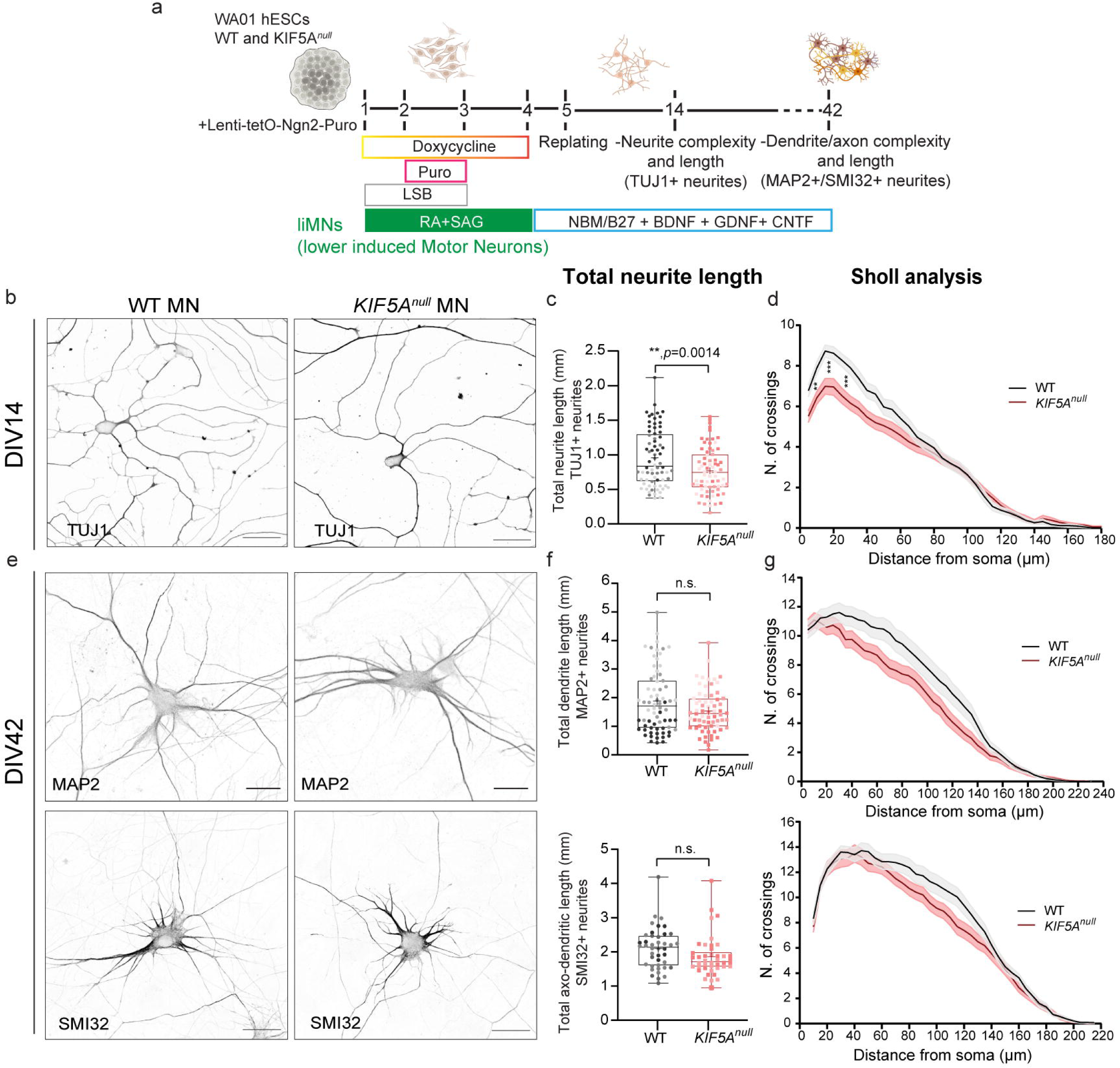
*KIF5A^null^* motor neurons display early defects in proximal neurite outgrowth in human motor neurons. (A) Diagram of experimental strategy to characterize neurite morphology at DIV14 and DIV42 in WT and *KIF5A^null^* lower MNs (B) Representative images of WT and *KIF5A^null^* MNs fixed at DIV14 and stained for the neurite marker TUJ1. Quantification of total TUJ1-positive neurite length (C) and Sholl analysis (D) of TUJ1-positive neurites in *KIF5A^null^* MNs at DIV14 in comparison to WT (n=3). (E) Representative images of WT and *KIF5A^null^* MNs fixed at DIV42 and stained for the dendrite marker MAP2 (top) and axo-dendritic marker SMI32 (bottom). Quantification of MAP2-positive dendrite (top) and SMI32-positive neurite (bottom) length (F) and Sholl analysis (G) of MAP2-positive dendrite (top) and *KIF5A^null^* MNs MNs at DIV42 in comparison to WT. (n=3 for MAP2, and n=2 for SMI32). Scale bars = 40 µm in (B) and 50 µm in (E). Data in (C) and (F) was assessed using a t-test with significance set to p < 0.05, and results are presented in boxplots with min/max whiskers, cross showing mean, and central bar is median. ** p < 0.01 Dots represent individual neurons, and each independent replicate is represented by different colors. Lines in Sholl analysis (D, G) represent sample means and shading represents the SEM, data was assessed with two-way ANOVA on linear mixed model, with replicates as random factor, post-hoc Tukey’s multiple comparison test was used to compare genotypes and rows considered distances. 10 µm: ** p < 0.01; 20 and 30µm: *** p < 0.001.

### Loss of KIF5A in human motor neurons leads to impaired axonal regrowth upon injury

Considering the importance of Kinesin-1-mediated transport in preserving the axonal structure^4,13,36–38^ we studied the role of KIF5A in axonal regeneration in motor neurons after injury. To that end, we cultured WT and KIF5A^null^ human motor neurons in compartmentalized microfluidic devices in which neurons extend axons through long (350µm) microgrooves^39^ and performed axotomy using vacuum aspiration at DIV14 or DIV42 (Figure 3a). Loss of KIF5A in motor neurons resulted in a substantially smaller number of recovered axons across all the time points measured post-injury both in DIV14 (Figure 3b-c) and DIV42 (Figure 3e-f) old neurons, with a significant reduced axonal regrowth at 3 days post-injury (Figure 3d,g). These results position KIF5A as an essential player in promoting axonal regeneration upon injury in human motor neurons at different time points during development.

**Figure 3.**
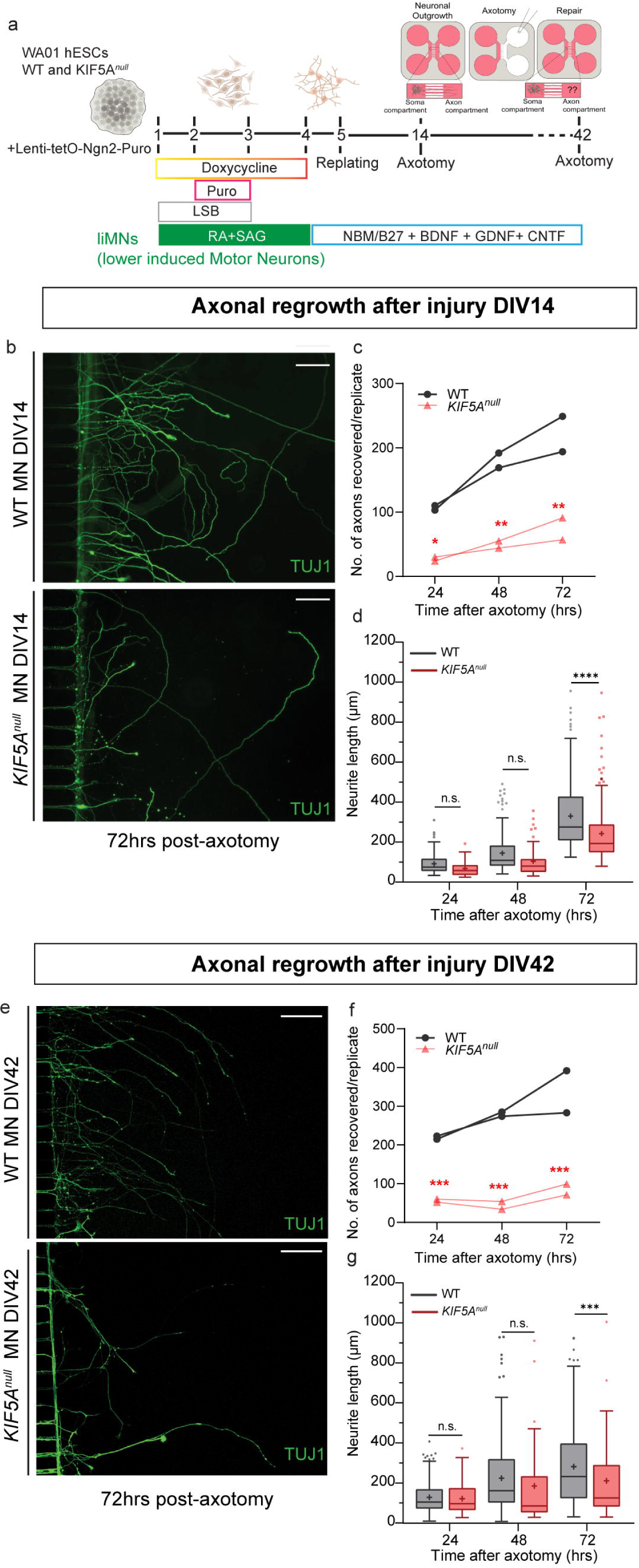
KIF5A depletion leads to axonal regrowth defects in human motor neurons at DIV14 and DIV42. (A) Diagram of experimental strategy to characterize neurite regrowth after injury at DIV14 and DIV42 in WT and *KIF5A^null^*MNs (B) Representative images of DIV14 liMNs in microfluidic devices 72hr post axotomy, stained for TUJ1 (green) (C) Quantification of number of recovered TUJ1+ axons at 24, 48 and 72hrs after axotomy in WT and *KIF5A^null^*MNs at DIV14 (D) Measurement of length of recovered axons at 24, 48 and 72hrs after axotomy after injury in WT and *KIF5A^null^* MNs at DIV14 (n=2). (E) Representative images of DIV42 MNs in microfluidic devices 72hr post axotomy, stained for TUJ1 (green) (F) Quantification of number of recovered TUJ1+axons at 24, 48 and 72hrs after axotomy in WT and *KIF5A^null^* MNs at DIV42 (G) Measurement of length of recovered axons at 24, 48 and 72hrs after axotomy after injury in WT and *KIF5A^null^* MNs at DIV42 (n=2). Scale bars = 100µm. Lines in line graphs (C) and (F) show mean, each line represents one replicate. Data was assessed using a Two-Way ANOVA with Sidak’s multiple comparison test with significance set to p < 0.05 in (C-G). * p < 0.05, ** p < 0.01, ***, p < 0.001, and **** p < 0.001. Results are presented in Tukey boxplots in (D,G) with cross showing mean and central bar median.

### KIF5A loss affect mitochondrial transport dynamics, but not neurofilament axonal trafficking in human motor neurons

Anterograde transport of mitochondria, one of the most frequently studied neuronal cargos transported by KIF5 motors, is crucial for ensuring neuronal integrity^24,40–43^. Impaired trafficking of these organelles is a common pathological feature in many neurodegenerative disorders, including ALS^2,13,36,38,44^. To assess whether KIF5A loss affects axonal transport in human motor neurons, we used live-cell imaging of mitochondria labeled with MitoTracker-Red in microfluidic devices at DIV14 and DIV42 (Figure 4 a,b). Motile mitochondria trafficked to a similar extent in both directions in WT motor neurons at DIV14 and DIV42 (Figure 4c-d). These organelles have been reported to become progressively less motile in maturing neuronal networks *in vivo*^45,46^. Most mitochondria were static both at DIV14 (73%) and DIV42 (71% static) in WT human motor neurons (Figure 4d) in line with previous reports in human neurons ^47^. At DIV14, no changes were observed in mitochondrial trafficking in KIF5A deficient motor neurons compared to WT (Figure 4c-e, Suppl. Figure 3a). However, at DIV42, *KIF5A*^null^ motor neurons showed a reduction in anterograde transport and an increased percentage of static mitochondria (35% anterogradely moving and 87% static mitochondria in *KIF5A*^null^ neurons *vs* 48% and 71%, respectively in WT neurons) (Figure 4d), accompanied by a reduction in average anterograde speed (0.48 µm/second *vs* 0.62 µm/second in WT) (Figure 4e, Suppl. Figure 3b).

**Figure 4.**
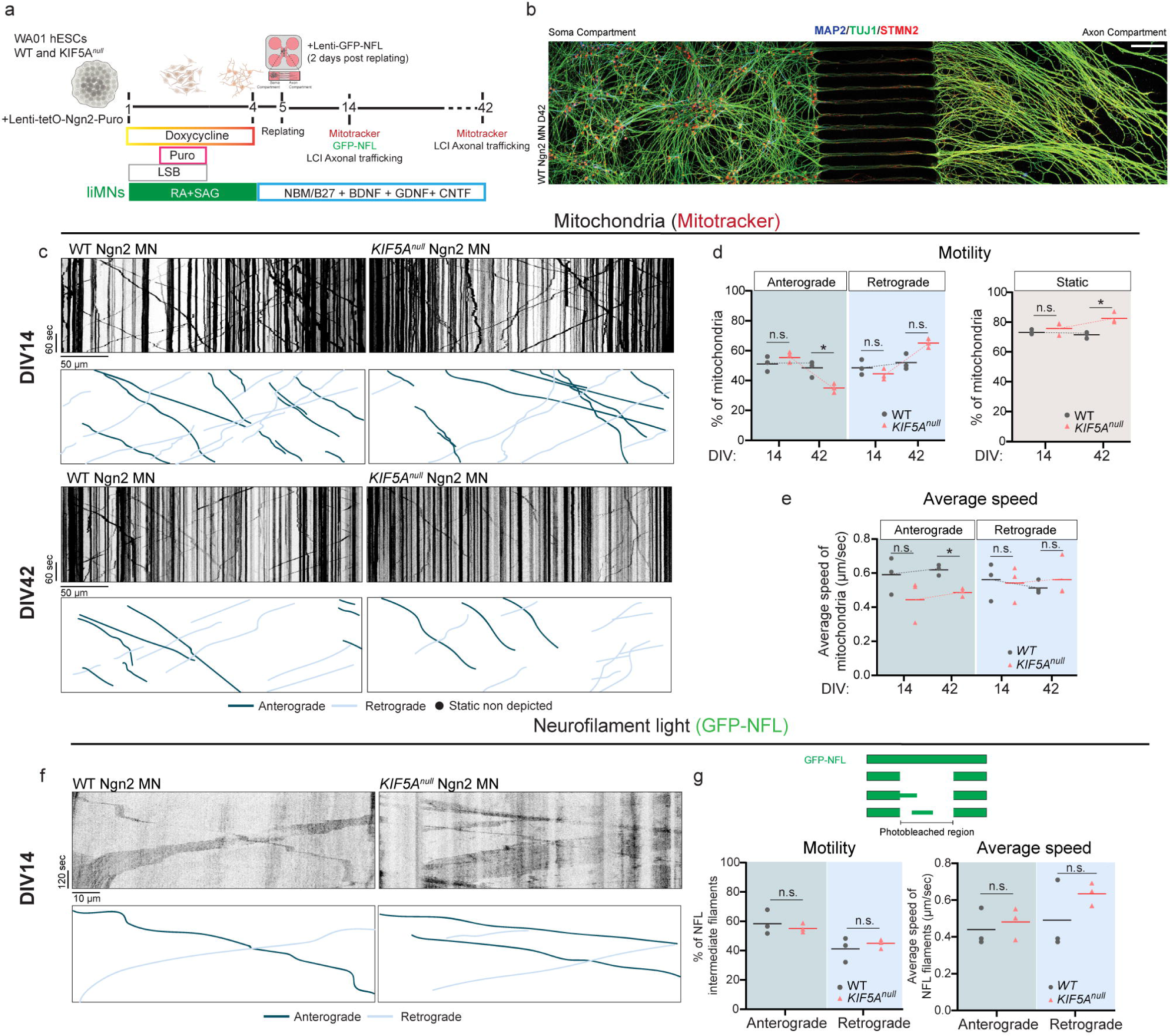
Anterograde axonal mitochondria trafficking is altered at DIV42, while NEFL trafficking is unaffected in KIF5a^null^ motor neurons. (A) Diagram of experimental strategy to study axonal mitochondria and neurofilament-light transport in WT and *KIF5A^null^* MNs at DIV14 and DIV42, respectively. (B) WT MNs cultured in compartmentalized microfluidic devices at DIV42 and stained for MAP2 (blue), β-III-Tubulin/TUJ1 (green) and STMN2 (red). (C) Representative kymographs of mitochondria labeled with Mitotracker-Red in WT and *KIF5A^null^* MNs at DIV14 (top) and DIV42 (bottom) and traces representing motile mitochondria, color coded for directionality. (D) Percentage of mitochondria fractioned into anterograde, retrograde (left panel) and static events (right panel) in WT and *KIF5A^null^* MNs at DIV14 and DIV42. (E) Average speed of anterogradely and retrogradely trafficked mitochondria in WT and *KIF5A^null^* MNs at DIV14 and DIV42 (n=3 independent replicates for DIV14 and DIV42). (F) Representative kymographs of GFP-NFL neurofilament light polymer dynamics in WT and *KIF5A^null^* MNs at DIV14 and traces representing motile Neurofilament-Light, color coded for directionality. (G) Schematic diagram of photobleaching technique (adapted from^58^) to study Neurofilament Light transport. GFP tagged Neurofilaments (GPP-NFL) are tracked through movement over photobleached gaps. (H) Percentage of Neurofilament Light trafficked polymers fractioned into anterograde, retrograde in WT and *KIF5A^null^* MNs at DIV14 (E) Average speed of anterogradely and retrogradely trafficked Neurofilament light filaments in WT and *KIF5A^null^*MNs at DIV14 (n=3). Bars represent median and dots are independent replicates. Data was assessed using a Two-Way ANOVA with Tukey’s multiple comparison with significance set to p < 0.05. Scale bars are indicated in Figure. Total time (vertical length) of kymographs are 5 minutes for Mitochondria and 10 minutes for Neurofilament-Light. * p < 0.05.

Neurofilaments (NFs) are transported along axons in a rapid, infrequent, intermittent, and bidirectional manner^48–50^. Abnormal NF accumulation, a hallmark of motor and peripheral neuropathies, has been proposed to be a consequence of disruptions in their transport by motor proteins^51–55^. KIF5 proteins, particularly KIF5A, are proposed as the anterograde motors for these polymers based on studies from *Kif5a* knockout mice ^56,57^. To examine the role of KIF5A in NF transport in human neurons, we imaged Neurofilament-Light (NFL) transport at DIV14 by expressing GFP-tagged NFL in KIF5A*^null^* and WT motor neurons in microfluidic devices (Figure 4a). Human motor neurons had no naturally occurring gaps or discontinuities in their neurofilament arrays to enable imaging of movement, unlike rodent cortical neurons^58,59^. Thus, we opted for a photobleaching strategy as previously described^59^ (Figure 4g, Suppl. Figure 3c). Neurofilament-Light polymers moved bidirectionally with 60% traveling in the anterogradely and 40% retrogradely, with a slightly higher average speed in the retrograde fraction in wild-type motor neurons (Figure 4f,g, Suppl. Figure 3d-e). No significant changes in neurofilament motility or average velocity in both directions were observed in motor neurons lacking KIF5A (Figure 4f,g, Suppl. Figure 3d-e). Hence, these data indicate that KIF5A is dispensable for Neurofilament-Light Chain transport in human motor neurons. We did not observe Neurofilament-medium trafficking at DIV14 or DIV42 in our motor neurons cultures nor NFL transport in older cultures (DIV42) even after photobleaching (Suppl. Figure 3c). Overall, our results show that loss KIF5A in human motor neurons leads to a time-dependent disruption of anterograde axonal mitochondrial transport, without affecting Neurofilament light trafficking.

### Axonal SFPQ granule anterograde transport is disrupted in mature *KIF5A^null^* motor neurons

Lastly, we investigated the effect of KIF5A loss on transport of ribonucleoprotein (RNP) granules, another cargo essential for maintenance of axonal function^4,60–64^. RNP granules are membraneless condensates composed of mRNA and RNA-binding proteins (RBPs) that travel from the cell body towards distal sites in axons and dendrites for local protein synthesis. Kinesin-1 motors have been previously reported to associate with RNP granules ^26,65^, especially, in the case of KIF5A, with those associated with the RBP splicing factor proline/glutamine–rich (SFPQ)^66^. SFPQ-RNA granule transport dynamics were visualized by expressing a Halo-tagged SFPQ in WT and *KIF5A*^null^ motor neurons at DIV14 and DIV42 in microfluidic chambers (Figure 5a). In DIV14 WT neurons, most RNA granules were motile (31% stationary), with a higher percentage moving in the retrograde direction (77%) and faster anterograde than retrograde speed (Figure 5b,d,e). SFPQ granule dynamics in DIV14 *KIF5A*^null^ motor neurons were not significantly different from those in WT motor neurons (Figure 5b,d,e, Suppl. Figure 4a). At DIV42, average bidirectional velocities were also similar between WT and KIF5a^null^ motor neurons (Figure5c,e, Suppl. Figure 4b). But, while the percentage of anterogradely moving granules almost doubled to 42% in WT motor neurons, the percentage in KIF5A mutant neurons remained at 20% (Figure 5c,d,). Together, our findings show that KIF5A becomes rate-limiting for anterograde transport of SFPQ-containing RNA granules in DIV42 neurons.

**Figure 5.**
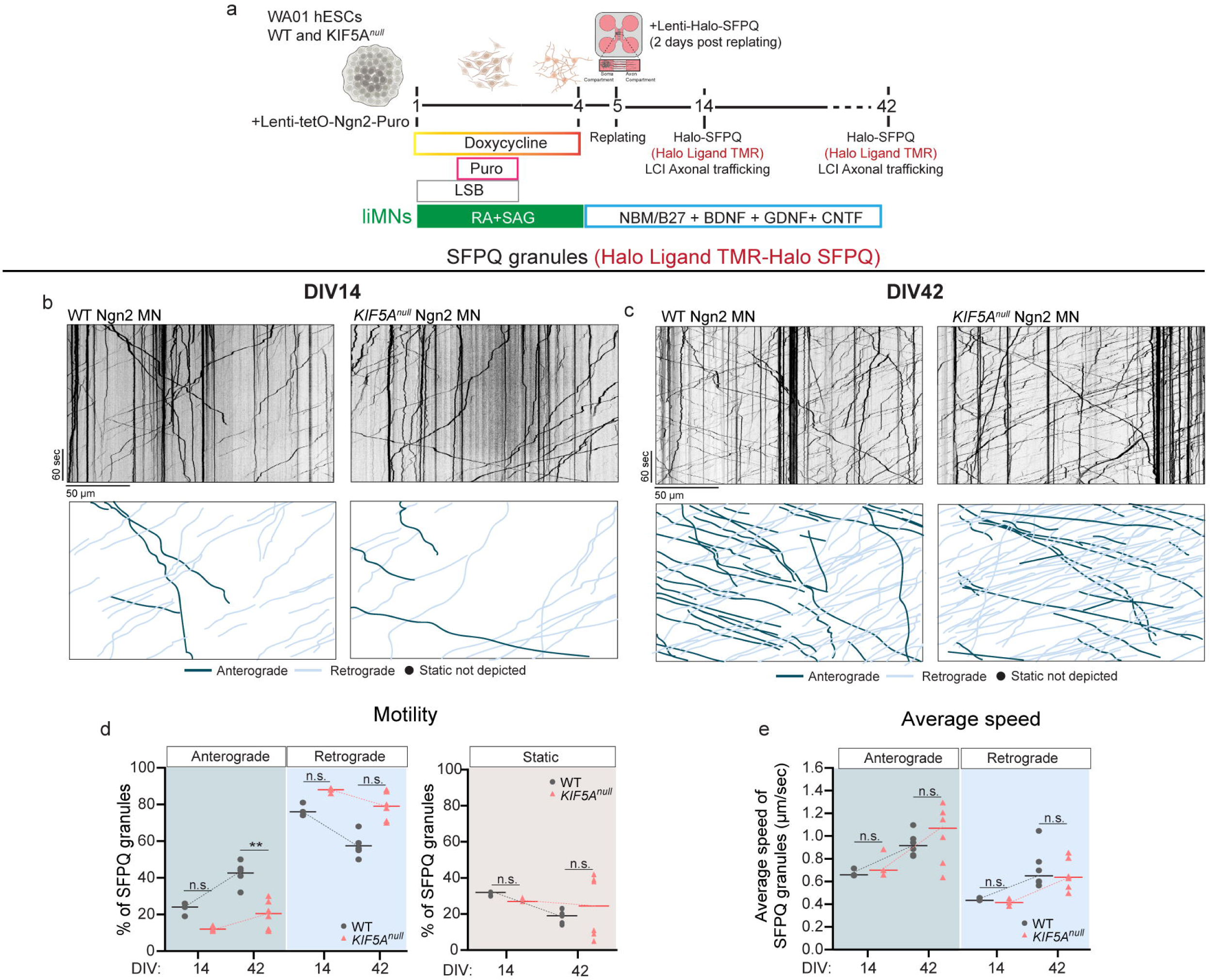
KIF5A loss affects anterograde axonal trafficking of SFPQ granules in motor neurons at DIV42. (A) Diagram of experimental strategy to study axonal Halo-SFPQ granule transport in WT and *KIF5A^null^* MNs at DIV14 and DIV42, respectively. (B) Representative kymographs of Halo-SFPQ granules labeled with Halo ligand TMR in WT and *KIF5A^null^* MNs at DIV14 and DIV42 (C) and traces representing motile mitochondria, color coded for directionality. (D) Percentage of Halo-SFPQ fractioned into anterograde, retrograde (left panel) and static events (right panel) in WT and *KIF5A^null^* MNs at DIV14 and DIV42. (E) Average speed of anterogradely and retrogradely trafficked Halo-SFPQ granules in WT and *KIF5A^null^* MNs at DIV14 and DIV42 (n=3 independent replicates for DIV14 and n=6 for DIV42). Bars represent median and dots are independent replicates. Data was assessed using a Two-Way ANOVA with Tukey’s multiple comparison with significance set to p < 0.05. Scale bars are indicated in Figure. Total time (vertical length) of kymographs are 5 minutes. ** p < 0.01. Detailed information per dataset (average, SD, n and detailed statistics) are available in Supplementary Table 1.

## Discussion

In this study, we examined the impact of KIF5A deficiency on neurite outgrowth under homeostatic and injury conditions as well as its effects on the axonal transport of various cargoes at two developmental stages, for the first time in human motor neurons. Loss of KIF5A led to a reduction in neurite complexity during early developmental stages and caused significant impairments in axonal repair capacity throughout development. KIF5A deficiency diminished axonal mitochondrial motility and, more substantially, transport of SFPQ-associated RNA granules, but did not affect neurofilament transport.

KIF5 paralogs follow a similar expression pattern during CNS development in rodents^26,27^ Publicly available datasets from human brain postmortem tissue obtained at different stages of development (BrainSpan, Allen Brain Atlas) indicated that KIF5A and KIF5C expression was most pronounced postnatally and during adulthood, with a distinctive increase of KIF5A, but not KIF5C, expression in the motor cortex (Figure 1A). In human stem cell-derived cortical and motor neurons, we observed a similar pattern of differential regulation of the KIF5 paralogs, with KIF5A expression increasing and KIF5B/C decreasing over time (Figure 1B). While some reports have found partial correlation with human postnatal and even adult stages^34^, stem-cell derived Ngn2 neuronal models do not fully recapitulate the identity of their *in vivo* counterparts^67–69^. Still, the distinctive regulation patterns we observe suggest unique functions for each KIF5 paralog in the developing and aging human brain. Moreover, the non-overlapping expression of KIF5 paralogs in humans might increase the susceptibility to effects of mutations in aging and developmental processes. For instance, the restricted expression of KIF5A in adulthood might make it more vulnerable to mutations related with late-onset neurological disorders. Such restricted temporal expression in combination with a regional specificity^70–72^ may explain why KIF5A is the only KIF5 paralog found to be mutated in adult-onset motor neuron disorders (e.g. ALS).

Here, we find that KIF5A regulates neurite outgrowth at early stages in human motor neuron development but becomes dispensable over time (Figure 3). KIF5 motors are one of the main components accumulating at axonal initiation sites in developing mammalian neurons^73,74^. In *Drosophila*, the single KIF5 ortholog (KHC) transiently drives initial axonal growth through cross-linking with MTs and promoting MT sliding, but this process is suppressed as neurons mature^75^. Hence, KIF5 appears to have a specific function early in axonal outgrowth, that is no longer necessary in mature neurons. Since the microtubule-binding sites on the KHC domain are conserved from *Drosophila* to humans, such a mechanism might also operate in human neurons, and could explain the observed deficits in neurite outgrowth at early, but not later stages.

Motor neurons developed normally in mice lacking *Kif5a*^76^ and retinal ganglion axon outgrowth was unaffected upon *Kif5a* loss in zebrafish^77^. However, mouse spinal motor neurons obtained from the same mice displayed reduced neurite outgrowth in culture^76^ and transient knockdown of KIF5 paralogs resulted in partially reduced neurite length in mouse hippocampal neurons *in vitro*^78^. Hence, two *in vivo* studies showed no impairments upon KIF5A loss, while two *in vitro* studies did. The collective evidence suggests that the specific KIF5A contribution to early axonal outgrowth might ultimately not be rate limiting for neuronal development *in vivo*.

At the systems level, genetic models of KIF5A dysfunction result in severe motor deficits. In *Drosophila*, expression of the Exon 27 skipped ALS mutant KIF5A reduced synaptic transmission at the neuromuscular junction (NMJ), altered NMJ morphology and affected locomotion^79^. *Kif5a* knockout mice died soon after birth due to a respiratory failure while postnatal inactivation of *Kif5a* in neurons resulted in loss of large caliber axons and hindlimb paralysis^56^. These findings suggest an essential function for *Kif5a* in motor neurons. Hence, while KIF5A expression may not be essential or at least rate limiting during brain development, the protein has an essential peri-/postnatal function in motor neurons.

At the cellular level, we find KIF5A to be essential in promoting axonal regrowth after injury in human motor axons *in vitro* (Figure 4). In line with this, *KIF5A* orthologs were shown to regulate axonal regeneration *in vivo* in *Drosophila* and zebrafish^80,81^. Additionally, knock-in of an ALS-associated mutation in the endogenous *Kif5a* mouse locus led to impaired axonal regeneration and subsequent recovery of motor units in response to injury and aging^82^. Together, these collective data suggest that impaired axonal repair underlies motor neuron dysfunction upon KIF5A disruption.

Our study shows that KIF5A depletion reduced axonal anterograde mitochondrial motility and to a greater degree SFPQ RNA granule anterograde transport in human motor neurons (Figure 4-5). The axonal mitochondrial transport speeds observed here are in the range of previously reported values (summarized in^83^) also in microfluidic devices^84^. These results are in agreement with previous studies that reported KIF5A orthologs as regulators of axonal transport of mitochondria in zebrafish^43^ and in primary motor neurons^76^. In line with the zebrafish study, we detected reduced motility and a specific reduction in the anterograde but not retrograde speed of axonal mitochondria in *KIF5A^null^* neurons (Figure 4c-e). Furthermore, our data on SFPQ RNA granule trafficking are in line with a study in dorsal root ganglion neurons showing that expression of Kif5a binding-deficient SFPQ resulted in 50% reduced anterograde transport of SFPQ granules in axons^66^. Moreover, in both studies, anterograde speed of RNA granules was unaffected. All in all, our data highlights the conserved role of KIF5A in anterograde axonal transport of these two cargoes. Notably, impairments in anterograde transport of mitochondria and SFPQ granules are evident only at DIV42 in human motor neurons, suggesting that increased KIF5A expression at later stages may underlie cargo specificity.

The anterograde transport of mitochondria and SFPQ RNA granule is not entirely blocked following KIF5A depletion (Figure 4-5), as other Kinesin motors may compensate for the absence of KIF5A. In fact, KIF5B was initially thought to be the motor protein responsible for anterograde mitochondrial transport^40,85,86^. Thus, even if KIF5B expression is reduced by half at DIV42 in motor neurons (Figure 1c-d), it could still be responsible for the residual mitochondria motility we observe in these neurons. Similarly, in the study where SFPQ was found to be selectively binding to KIF5A among KIF5 motors^66^, members of the Kinesin-3 family (KIF1) were also identified in the coprecipitates of SFPQ. Interestingly, in zebrafish, concurrent loss of the KIF1B and KIF5A orthologs exacerbates axonal degeneration through transport of an unknown nonmitochondrial cargo shared by both motors^43^. Given that mitochondria are a conserved cargo of KIF5A across species, RNA granules might too be a common conserved cargo for both KIF5A and KIF1B in human motor neurons. Overall, our work provides additional evidence highlighting the complexity of Kinesin redundancy and cargo specificity in human motor neurons.

The mouse ortholog of *KIF5A* was found to be the principal motor responsible for neurofilament medium transport in primary cultures^57^. *In vivo*, the evidence is less direct: conditional knockout of *Kif5a* showed a specific increase in neurofilament light and heavy levels in the cell bodies of DRGs after sciatic nerve ligation^56^. In contrast, our findings indicate that KIF5A is not essential for neurofilament light transport in human motor neurons (Figure 4f-h). Similarly, a separate study where Kif5a was conditionally deleted in neurons *in vivo,* reported no accumulation of neurofilaments in the CNS^87^ and mutations in the cargo binding region of the *Kif5a* zebrafish ortholog showed no defects in axonal neurofilament localization^88^ Notably, in DRG mouse cultures knockdown of dynein/dynactin and expression of an N-terminal *Kif5c* mutant also inhibited anterograde and retrograde transport of neurofilament medium^57^ Our transport parameters are in line with those reported in previous studies where photobleaching of neurofilaments was carried out^59^. However, we are unable to draw any conclusions on Neurofilament light transport at later stages or Neurofilament medium altogether based on the technical limitations associated with imaging movement of these polymers in neuronal cultures^58^. Still, these data collectively suggest that KIF5A is not the exclusive anterograde motor in neurons.

The axonal transport deficits reported here might contribute to the impairments in axonal repair in the absence of KIF5A. In mice, Kif5a transport was found to be the most significantly affected process in retinal axons undergoing repair, and retinal ganglion cells lacking *Kif5a* showed defective mitochondrial and RNP transport upon injury^89^. These findings suggest that KIF5A regulates axonal repair through transport of cargoes essential for maintaining axonal integrity. Increasing evidence points to mitochondrial recruitment in regenerating axons as one of the critical factors that promotes axonal repair (reviewed in ^90,91^). For instance, overexpression of the mitochondrial protein ARMCX1 enhances axonal mitochondrial transport and facilitates axon regeneration after injury^92^. In addition, RNP granule transport might be important for recruitment of locally translated RNAs necessary for injury responses. In *C. elegans*, ribosomes are recruited to regenerating axonal tips and local translation is induced shortly after injury^93,94^, and SFPQ granules transport mRNAs that are locally translated and are important for axonal survival in mouse DRG neurons^66,95–100^. *Bclw*, one of the SFPQ-transported RNAs, is locally translated^96^ and its mimetic peptide prevents axonal degeneration in DRG neurons expressing mutant Kif5a^66^. Although the human RNAs that are trafficked by SFPQ remain unknown, we anticipate similar mechanisms that promote axonal survival responses in human motor neurons. We postulate that deficits in the axonal transport of mitochondria and SFPQ granules and potentially other cargoes explain motor axon dysfunction.

Finally, we speculate that axonal transport and repair could similarly be affected in motor neurons in the context of ALS pathogenic mutations. Initially ALS variants in KIF5A were predicted to be loss-of-function^14^ but recent studies demonstrate a gain of function mechanism in which mis-splicing results in a novel C-terminal domain that prevents autoinhibition and hence renders the kinesin hyperactive^31,32,101,102^. Yet, the sequence changes in the C-terminal domain probably also impair cargo binding and thus impair KIF5A-dependent transport, e.g., axonal SFPQ RNA granules and mitochondria transport as demonstrated in the current study. Indeed, mutations in the C-terminal tail of Kif5a in zebrafish, significantly impair axonal localization of mitochondria^88^. Impaired transport of these two cargoes and potentially others, may reduce axonal stability, result in neuromuscular junction (NMJ) dysfunction, and eventually denervation. In fact, expression of a mis-spliced KIF5A mutant in Drosophila^79^ or knock-in of an ALS mutation in the endogenous *Kif5a* mouse locus^82^ resulted in NMJ instability, locomotor alterations and impaired motor recovery after injury. Considering that mutant KIF5A forms hybrid dimers with WT KIF5A^32^, it might also form hybrid dimers with other KIF5 motors^24^ and consequently impair their functions too. Such chimeric mutant/wild-type KIF5 complexes might prevent compensatory or adaptive processes to occur. Thus, these ALS mutant motor neurons might, in contrast to *KIF5A^null^*motor neurons, additionally show defects in neurofilament transport and axonal outgrowth. In closing, disturbed axonal transport of SFPQ and mitochondria provides a plausible mechanism for mutant KIF5A-mediated axonal transport defects in ALS pathogenesis.

## Supporting information

Supplemental mitotracker videos

Supplemental neurofilament videos

Supplemental SFPQ videos

## Supplemental Figures

**Supplemental Figure 1.**
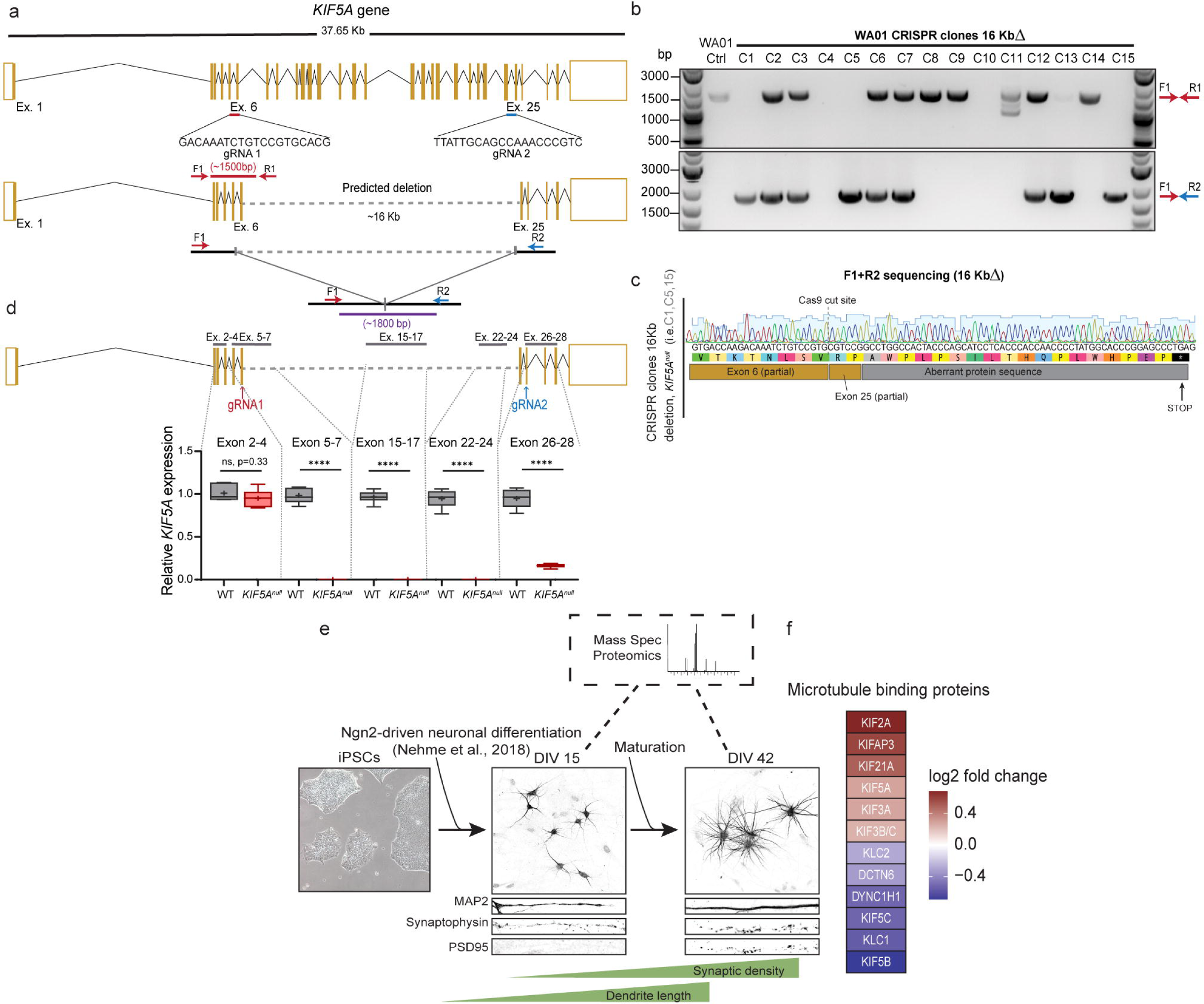
Generation of *KIF5A^null^* motor neurons via CRISPR/Cas9 editing and opposite regulation of functionally related proteins in Ngn2 neuron developmental proteome. (A) Predicted 16kb deletion in *KIF5A* locus. Primer pairs were designed to flank the 16Kb deletion region to confirm the presence or absence of mutations. (B) Genotyping PCR of WA01 hESCs clones at the indicated sites representing a summary of mutations identified within the KIF5A allele as the result of CRISPR editing. (C) Representative sequencing results for the KIF5A editing of selected CRISPR hESC clones. (D) Examination of KIF5A mRNA in differentiated motor neurons from the selected clone, spanning different regions of the gene, upstream, downstream and within deletion (n=3 technical replicates). Data are presented in Tukey boxplots with cross showing mean and central bar median and analyzed with a t-test with significance set to p < 0.05. **** p < 0.0001. (E) Schematic experimental approach: iPSCs from three healthy individuals were differentiated into iNeurons using NGN2 expression in combination with dual SMAD and Wnt inhibition^34^ iNeurons were harvested 15 (DIV15) and 42 days (DIV42) after induction, for at least two inductions per time point and protein levels were measured using Mass Spectrometry proteomics. Supplemental table 2 contains the complete list of DEA proteins. (F) Opposite regulation within protein class “Microtubule binding motor protein”. Color coding indicates log2 fold change at DIV42 compared to DIV15. Supplemental table 3 contains the complete list of panther DB protein family classifications.

**Supplemental Figure 2.**
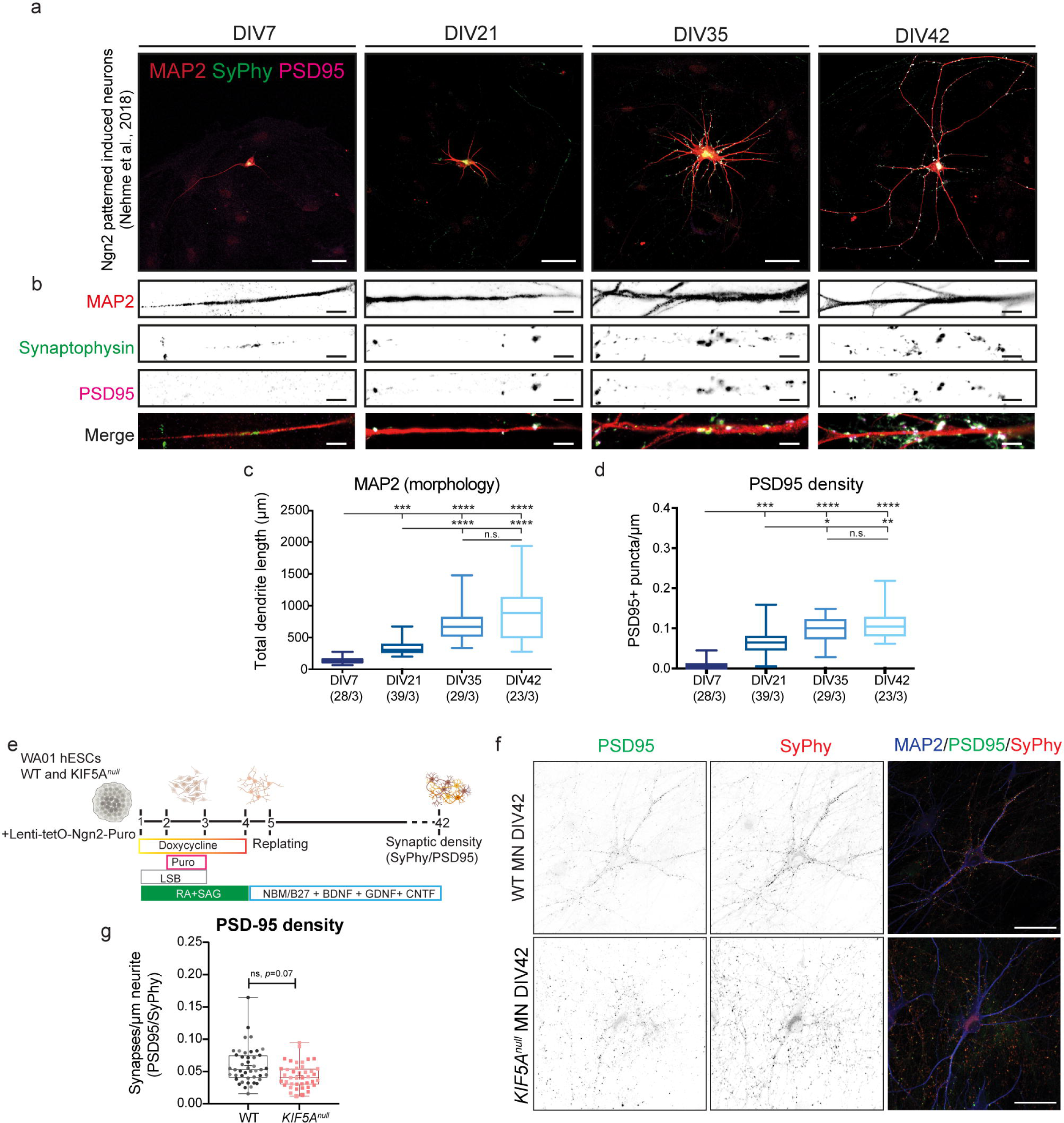
Absence of KIF5A does not result in defects in synaptic density at DIV42. (A) Examples of DIV7, DIV21, DIV35 and DIV42 stained for MAP2 (red), Synaptophysin (green) and PSD95 (magenta). Scale bars=50µm. (B) Dendrite zooms of neurons in (A), scale bars=5µm. Quantification of dendrite length (C) and PSD95 puncta density (D). Results are shown as min-to-max boxplots, sample sizes are indicted below each graph (n/N). (E) Diagram of experimental strategy to characterize neurite regrowth after injury at DIV14 and DIV42 in WT and *KIF5A^null^* MNs (F) Representative images of in WT and *KIF5A^null^* MNs at DIV42 stained for Synaptophysin (red), PSD-95 (green) and MAP2 (magenta) and quantified (G) for density of PSD-95/SyPhy puncta/µm. Scale bars=50 µm. (N=2 technical replicates). Results are presented in min-to-max boxplots, and central bar is median. Dots represent individual neurons, and each independent replicate is represented by different colors. Data was assessed with Kruskal-Wallis with Dunn’s multiple comparison test in (C,D) and a t-test in (G), significance set to p < 0.05. * p < 0.05, **p < 0. 01, *** p < 0.001, and **** p < 0.0001.

**Supplemental figure 3.**
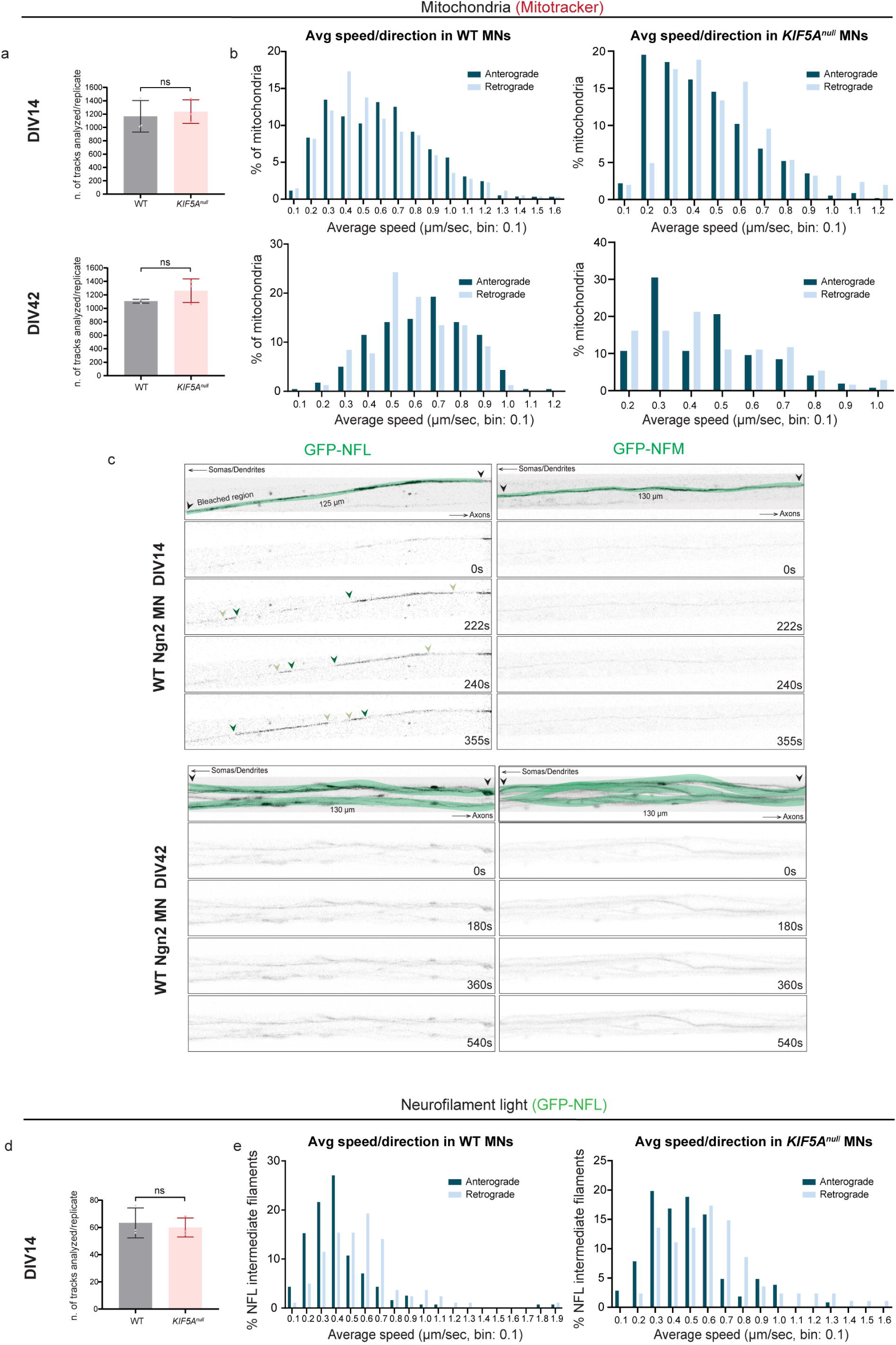
Dynamics of axonal mitochondria and NFL transport in WT and *KIF5A^null^* motor neurons at DIV14/DIV42 and DIV14 only, respectively. (A) Quantification of number of mitochondria tracks analyzed for each experimental replicate for trafficking analysis in WT and KIF5A*^null^* MNs at DIV14 (top) and DIV42 (bottom). (B) Distribution of mitochondria velocity in the anterograde (dark blue) or retrograde (light blue) direction in WT and KIF5A*^null^* MNs at DIV14 (top) and DIV42 (bottom). (C) Representative still images of time-lapse movies showing the fluorescence recovery of axonal neurofilaments containing GFP-tagged NFL (left panel) or NFM (right panel) at DIV14 (top) or DIV42 (bottom) in WT MNs. Movement of single neurofilament polymers through photobleached gaps (green line) in the axonal neurofilament array only occurs in MNs expressing GFP-tagged NFL at DIV14 (top left panel). The leading end of the moving filament is indicated by a dark green arrow, and the trailing end by a light green arrow. Time points of each image are indicated at the bottom right of each frame. (D) Quantification of number of GFP-NFL neurofilament tracks analyzed for each experimental replicate for trafficking analysis in WT and KIF5A*^null^* MNs at DIV14. (E) Distribution of NFL velocity in the anterograde (dark blue) or retrograde (light blue) direction in WT and KIF5A*^null^* MNs at DIV14.

**Supplemental Figure 4.**
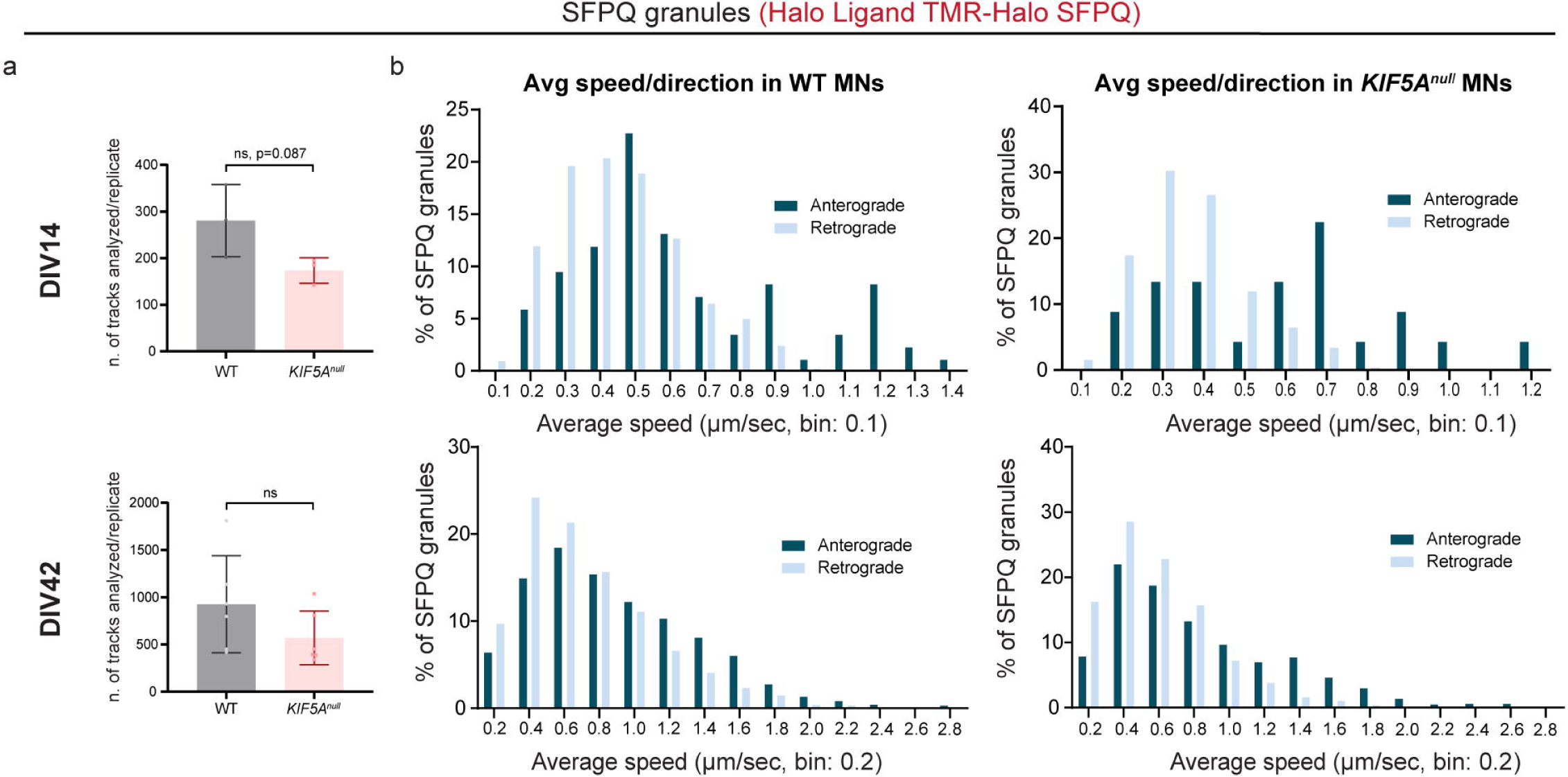
Dynamics of axonal SFPQ granule transport in WT and *KIF5A^null^* motor neurons at DIV14/ DIV42. (A) Quantification of number of Halo-SFPQ tracks analyzed for each experimental replicate for trafficking analysis in WT and KIF5A*^null^* MNs at DIV14 (top) and DIV42 (bottom). (B) Distribution of SFPQ granule velocity in the anterograde (dark blue) or retrograde (light blue) direction in WT and KIF5A*^null^* MNs at DIV14 (top) and DIV42 (bottom).

## Materials and Methods

### CRISPR Guide Design, validation, and generation of *KIF5A^null^* hESCs

KIF5A guide RNAs were designed using the following web resources: CHOPCHOP (https://chopchop.rc.fas.harvard.edu) from the Schier Lab. Guides were cloned into a vector containing the human U6 promotor (custom synthesis Broad Institute, Cambridge) followed by the cloning site available by cleavage with BbsI, as well as ampicillin resistance. To perform the cloning, all the gRNAs were modified before ordering. The following modifications were used in order to generate overhangs compatible with a BbsI sticky end: if the 5’ nucleotide of the sense strand was not a G, this nucleotide was removed and substituted with a G; for the reverse complement strand, the most 3’ nucleotide was removed and substituted with a C, while AAAC was added to the 5’ end. The resulting modified STMN2 gRNA sequences were used for Cas9 nuclease genome editing guide 1: 5’ GACAAATCTGTCCGTGCACG 3’ (Exon 6), guide 2: 5’ TTATTGCAGCCAAACCCGTC 3’ (Exon 25). Cloning was performed by first annealing and phosphorylating both the gRNAs in PCR tubes. 1 µL of both the strands at a concentration of 100 µM was added to 1 µL of T4 PNK (New England Biolabs), 1 µL of T4 ligation buffer and 6 µL of H2O. The tubes were placed in the thermocycler and incubated at 37°C for 30 mins, followed by 5 mins at 95°C and a slow ramp down to 25°C at a rate of 5°C/minute. The annealed oligos were subsequently diluted 1:100 and 2 µL was added to the ligation reaction containing 2 µL of the 100 µM pUC6 vector, 2 µL of NEB buffer 2.1, 1 µL of 10mM DTT, 1 µL of 10mM ATP, 1 µL of BbsI (New England Biolabs), 0.5 µL of T7 ligase (New England Biolabs) and 10.5 µL of H2O. This solution was incubated in a thermocycler with the following cycle, 37°C for 5 minutes followed by 21°C for 5 minutes, repeated a total of 6 times. The vectors were subsequently cloned in OneShot Top10 (ThermoFisher Scientific) cells and plated on LB-ampicilin agar plates and incubated overnight on 37°C. The vectors were isolated using the Qiagen MIDI-prep kit (Qiagen) and measured DNA concentration using the nanodrop. Proper cloning was verified by sequencing the vectors by Genewiz using the M13F (−21) primer.

Human embryonic stem cell transfection was performed using the Neon Transfection System (ThermoFisher Scientific) with the 100 µL kit (ThermoFisher Scientific). Prior to the transfection, stem cells were incubated in mTeSR1 containing 10µM Rock inhibitor for 1 hour. Cells were subsequently dissociated by adding accutase and incubating for 5 min at 37°C. Cells were counted using the Countess and resuspended in R medium at a concentration of 2,5*10^6^ cells/mL. The cell solution was then added to a tube containing 1 µg of each vector containing the guide and 1.5 µg of the pSpCas9n(BB)-2A-Puro (PX462) V2.0, a gift from Feng Zhang (Addgene). The electroporated cells were immediately released in pre-incubated 37°C mTeSR medium containing 10µM of Rock inhibitor in a 10-cm dish when transfected with the puromycin resistant vector. 24 hours after transfection with the Puromycin resistant vector, selection was started. Medium was aspirated and replaced with mTESR1 medium containing different concentrations of Puromycin: 1µg/µL, 2µg/µL and 4 µg/µL. After an additional 24 hours, the medium was aspirated and replaced with mTeSR1 medium. Cells were cultured for 10 days before colony picking the cells into a 24-well plate for expansion.

Genomic DNA was extracted from puromycin-selected colonies using the Qiagen DNeasy Blood and Tissue kit (Qiagen) and PCR screened to confirm the presence of the 16 Kb deletion in the STMN2 gene. PCR products were analyzed after electrophoresis on a 1% Agarose Gel. In brief, the targeted sequence was PCR amplified by a pair of primers external to the deletion, designed to produce a ∼1800 bp deletion-band in order to detect clones containing the large deletion. Sequences of the primers used are as follows: F1 5’ TCCAGAGGCAGATAGATGAG 3’, R1 5’ CTATTTGGGACACTGAAGCAG 3’, R2 5’ GCCAAGGCTTCAAATA 3’. Geneious software was used to determine the changes in the ORF (Suppl. Figure 1c).

### iPSC/hESC lines

Three iPSC lines from unrelated individuals with no diagnosed disease status were used for MS proteomics. The hESC line was used for CRISPR/Cas9 editing. Details for each line, such as source material and reprogramming method are listed in the table below. iPSCs/hESCs were routinely tested for mycoplasma contamination (every two-three months) and checked for bacteriostasis/fungistasis (daily).

| Cell line information | Reprogramming technique | Source | Identifier | Passages Received frozen stocks Upon NGN2 infection Upon induction |
| --- | --- | --- | --- | --- |
| GM25256. Male skin fibroblast, 30 yo healthy control, Asian. Episomal reprogramming. | Episomal | Coriell Institute |  | P40<br>P48<br>P50 |
| BIONi010-C-6. Male fibroblast, 15-19y, Black/African American. Episomal reprogramming. | Episomal | Bioneer | Biosamples ID: SAMEA4454009 | P60<br>P63<br>P65 |
| GM23973. Male skin fibroblast, 19 yo healthy control, Caucasian. Viral reprogramming | Viral | Coriell Institute,<sup>103</sup> |  | P40<br>P43<br>P45 |
| WA01 (H1) cell line, male, human embryonic stem cell line, derived from blastocyst stage, European, none reported disease. | NA | WiCell Research Institute.<sup>104</sup> | RRID: CVCL_9771 | P23<br>P25(for CRISPR editing)<br>P30<br>P32 |

#### Cell culture and differentiation of iPSCs/hESCs into cortical/motor neurons

Induced pluripotent stem cells and human embryonic stem cells were maintained in Essential E8 medium (Gibco #A1517001) on Matrigel-coated plates (VWR # BDAA256277), passaged using 1mM EDTA, AND maintained in 5% CO2 incubators at 37 °C. 10 μM ROCK inhibitor (Sigma, Y-27632) was added to the cultures for 24 h after replating to prevent cell death. hESCs and iPSCs were co-infected with TetO-Ngn2-Puro (Addgene #52047) and reverse tetracycline-controlled transactivator (rtTA) (Addgene #19780) and were plated at a density of 100,000 cells/cm^2^ with rock inhibitor Y27632 (Stemgent 04-0012) on Geltrex-coated culture plates. Infected iPSCs were subsequently expanded for a couple of passages (maximum of five) to obtain a large batch of iPSCs with the NGN2 cassette and to eliminate presence of active viral particles.

Cortical neuron differentiation was induced as described previously^34^, cells were grown in N2 medium (DMEM/F12 medium (Life Technologies # 10565018), supplemented with 200mM Glutamax (Life Technologies # 1157446), 20% Dextrose (Life Technologies # A2494001), 1% N2 supplement B (Stemcell Technologies # 07156) and 0.1% Pen/Strep) to which doxycycline hyclate (2µg/ml; Sigma # D9891) and dual SMAD inhibitors (100nM LDN193189 (Stemgent #04-0074), 10µM SB431542 (Tocris # 1614), 2µM XAV939 (Stemgent # 04-00046) were added. After 24 hours of induction, culture medium was replaced with the same components but including puromycin (Merck/Millipore # 540222) as a selection step (puromycin concentration required was determined separately for each iPSC line). This step was repeated for another 24 hours. The next day, medium was replaced by N2-supplemented culture medium with 10µM FUDR (Sigma # F0503) for another 24 hours, after which neurons were replated onto poly-ornithine (Sigma #P6407)/laminin (Sigma #L2020)-coated 6-well plates for proteomic (density of 300k neurons/well), Motor neuron differentiation was achieved using a modified strategy from the previously reported NGN2-driven reprogramming protocol coupled with activation of posteriorizing and ventralizing signaling pathways via addition of patterning factors, retinoic Acid (RA, 1 µM, Tocris) and smoothened agonist (SAG, 1 µM, StemCell Technologies) for three days^35^. On day 5, cells were dissociated, replated in either poly-ornithine (Sigma #P6407)/laminin (Sigma #L2020)-coated 6-well plates for Western Blot (density of 300k neurons/well), 12-well glass coverslips (25k-30k neurons/well) with or without glia feeder layers for immunocytochemistry and PDMS microfluidic devices on POL-coated glass bottom dishes (75k neurons/device) for transport assays.

### SNP analysis and CNV calling for hPSCs lines

Prior to induction of NGN2, hPSCs were subjected to SNP-array analysis to confirm absence of large-scale genomic aberrations. DNA was prepared using QIAGEN DNeasy Blood & Tissue kit and processed by the Infinium global screening array (Illumina). CNV calling was performed using the iPsychCNV package in R (http://biopsych.dk/iPsychCNV). CNVs larger than 500Kb (containing >100 SNPs) were called if the region was flagged by two analysis metrics, one based on the Log R ratio and one using B allele frequency. Subsequently, called CNVs are compared against gene lists compiled based on gene ontology (GO) terms for ‘brain development’, ‘synapse’, and ‘cytoskeleton’. If genes affected by an identified CNV also occur in this gene list, the sample was excluded.

### Constructs used in this study and lentiviral infection

The EGFP-NFL construct was generated from the human neurofilament light cDNA, obtained from the expression plasmid from Anthony Brown’s lab (Addgene #132589, ^105^, cloned into a pEGFP-C1 plasmid to label the NFL with EGFP at the N-terminus, and subsequently cloned to a Synapsin-promoter driven lentiviral construct. The lentiviral UbC-Halo-SFPQ construct was acquired from Addgene (#166946), which was generated and described previously by Rosalind Segal’s lab^66^. All constructs were sequence verified before production of viral particles. To visualize neurofilament light and SFPQ granule transport, lower induced human motor neurons in microfluidic devices were infected with EGFP-NFL or Halo-SFPQ lentiviral particles at DIV7. Motor neurons were imaged at DIV14 (7 days post infection) for neurofilament and SFPQ granules and additionally at DIV42 (35 days post infection) for SFPQ granules.

### Mass spectroscopy, MS proteomics data analysis

#### Mass spectroscopy

Cells were washed 2 times with ice cold PBS. 500uL protease inhibitor (PI) solution in PBS (complete EDTA-free PI tablets, Roche 05056489001) was added to each well. Cells were collected by gentle scraping and spun down for 5 minutes at 3000 rcf at 4C. Supernatant was removed, and the pellet was resuspended in 20uL loading buffer (4% SDS, 100mM Tris pH 6.8, 0.04% bromophenol blue, 200mM DTT, 20% glycerol, and PI in PBS). Samples were snap frozen and stored at −80C until further processed. An SDS–PAGE LC-MS/MS approach was used for protein identification as described previously (https://doi-org.vu-nl.idm.oclc.org/10.1007/978-1-4939-9662-9_11). In brief, after the sample was run into the gel, the protein-containing gel piece was excised and digested with trypsin/Lys-C mix (Promega). Peptides were analyzed by a data-independent acquisition (SWATH) method in a TripleTop 5600+ mass spectrometer (Sciex).

A spectral library was made from pooled samples of all three lines, one for each culture condition, collected at two different time points (DIV15 and DIV42). Additionally, a pooled sample glia cultured without neurons, collected at DIV15 and DIV42, was included. Spectral library samples were measured in DDA mode and analyzed using MaxQuant 1.6.3.4^106^ (. The Uniprot human reference proteome database (SwissProt + TrEMBL, version 2019-11) was used to annotate spectra. The minimum peptide length was set to 6, with at most two miss-cleavages allowed. Methionine oxidation and N-terminal acetylation were set as variable modifications with cysteine Propionamide set as fixed modification. For both peptide and protein identification a false discovery rate of 0.01 was set.

SWATH data were searched against the spectral library (peptides and proteins identified from DDA data by MaxQuant) using Spectronaut 13.7^107^ with default settings. The resulting abundance values and qualitative scores for each peptide in the spectral library were exported for further downstream analysis.

#### MS proteomics data analysis

MS proteomics data analyses were performed using the R language for statistical computing. MS-DAP 0.2.6.4 (https://github.com/ftwkoopmans/msdap;^108^) was used for the interpretation of data quality and differential expression analysis (DEA). While importing the Spectronaut data report, fragment group MS2 total peak areas without Spectronaut normalization were selected to represent peptide intensity values and both proteins from the MaxQuant contaminant database and iRT peptides were removed from the dataset.

In the statistical contrast, peptides observed with Spectronaut confidence score <= 0.01 in all samples of both time points were selected. MS-DAP’s mode-between normalization was then applied to this data subset and finally the limma eBayes statistical model was used for differential testing between the two time points^109^. The significance threshold was set at 1% FDR. All data visualizations and MS-DAP parameters are included in the MS-DAP report. For detection analysis, all proteins detected in all lines with at least two samples per line were included for each time point. Functional enrichment analysis was performed using g:GOSt in g:Profiler^110^ (https://biit.cs.ut.ee/gprofiler/gost Enriched GO terms were grouped using the cytoscape plugin ClueGO^111^. The SynGO database was used for filtering of SynGO annotated proteins, SynGO enrichment analysis and visualization of protein counts per SynGO term^112^ (www.syngoportal.org). For enrichment analysis with both g:Profiler and SynGO, the total DEA dataset was as custom background list. The PANTHER (Protein ANalysis THrough Evolutionary Relationships) functional classification system was used to group significantly regulated proteins into protein classes^113^ (http://www.pantherdb.org. The String Database^114^ (StringDB was used to visualize the interactions between downregulated synaptic organization proteins.

### Immunocytochemistry and morphological analysis

Neurons were fixed at different time points (indicated in figures) with 3.7% paraformaldehyde or 100% ice cold Methanol (PFA; Electron Microscopy Sciences) for synaptic stainings, then washed three times with PBS pH = 7.4. Cultures were permeabilized with 0.5% Triton X-100 (Thermo Fisher #T/3751/08), followed by a 30 min incubation in PBS containing 2% normal goat serum (NGS; Thermo Fisher #11540526) and 0.1% Triton X-100. Next, neurons were stained with primary antibodies overnight at 4°C. The following antibodies were used: chicken anti-MAP2 (1:300, Abcam Ab5392), Guinea pig anti-Synaptophysin 1 (1:1000, Synaptic Systems #101004), rabbit anti-Synapsin (1:300, E028), mouse anti-PSD95 (1:500, Abcam ab2723), mouse anti-βIII tubulin (1:300 Invitrogen MA1-19187), rabbit anti-KIF5A (1:100 Abcam ab5628), mouse anti-SFPQ (1:100 Abcam ab11825), rabbit anti-STMN2 (1:300 1:300 Abcam ab185956). After three washes with PBS, neurons were stained with secondary antibodies Alexa Fluor (1:1000; Invitrogen) and counterstained with DAPI (1:1000) for 1h at RT. Following three additional washes, coverslips were mounted on microscopic slides with Mowiol-DABCO or for microfluidic devices on glass-bottom dishes, were ready to be imaged. Images were acquired on a Nikon Ti-Eclipse microscope equipped with a confocal scanner model A1R+, using a 40X oil immersion objective (NA=1.3; Nikon) or a 60X oil immersion objective (NA=1.4; Nikon). Settings were optimized for each time point for each experiment but kept the same between cells and coverslips or microfluidic devices. SynD automated analysis software in MATLAB was used to quantify neuronal morphology and density of synaptic puncta^115^ MAP2 masks are drawn using thresholding and manual tracing to quantify dendrite length. A synapse mask is drawn with thresholding of the synapse marker; requiring putative synapses to be within the dendrite mask, above the mean synapse channel intensity by a manually set number of standard deviations, and above a manually set area size. The synapse mask is used to quantify synapse numbers and intensity. Synapse detection settings were optimized for each time point per culture.

#### Western blotting

For examination of KIF5 protein levels, motor neurons were lysed in RIPA buffer (150mM Sodium Chloride; 1% Triton X-100; 0.5% sodium deoxycholate; 0.1% SDS; 50 mM Tris pH 8.0) containing Halt protease and phosphatase inhibitors (Life Technologies 78441) and centrifuged at 12,000 RPM for 10 minutes at 4°C. Protein concentration was determined by a BCA assay (Thermo Scientific 23225) and 10-20 μg of total protein were separated by SDS-PAGE using a 4-20% gradient (Bio-Rad 4561094), transferred to polyvinylidene difluoride membranes (EMD Millipore IPFL00010) and probed with antibodies against GAPDH (1:2,000, mouse monoclonal, EMD Millipore MAB374) and KIF5A (1:500, rabbit polyclonal Abcam ab5628), KIF5B (1:1,000, rabbit polyclonal proteintech 21632-1-AP) and KIF5C (KIF5C) levels were normalized to GAPDH. LiCor software (Image Studios) was used to visualize and quantitate protein signal.

#### RNA Isolation and qRT-PCR

RNA was isolated from motor neuron lysates with RNA Qiagen RNeasy Plus Micro kit (Qiagen 74034) or Trizol (Invitrogen), in accordance with manufacturer’s recommendations and quantified spectrophotometrically (at 260 nm). Approximately 300ng of total RNA was used to synthesize cDNA using the iScript kit reverse transcriptase (Bio-Rad 1708891). The cDNA was then amplified using the SYBR Premix (iScript Advanced cDNA Synthesis Kit) using the CFX96 Touch Real-Time PCR Detection System StepOnePlus (Bio-Rad). Briefly, each 20 μl of reaction volume contained 5 μL of SYBR Green PCR Master Mix or 5 μL PrimeTime™ Gene Expression Master Mix, 0.5 μM of each primer, and UltraPure destilated water DNase and RNase free (Invitrogen). Genes were normalized to GAPDH (SYBR) expression and expressed relative to their respective control condition. All KIF5A exon junctions were analyzed for in three biological replicates with each have technical duplicates.

### Fabrication and use of microfluidic devices

PDMS-based microfluidic devices were generated based on a protocol described in detail by Yong et al., 2020 with slight modifications. Briefly, two high resolution chromium on glass photomasks based on designs from the Deppman Lab^39^ were designed and fabricated (Mask files available in Supplementary Material). These masks were used in I-line photolithography to pattern a two-layer SU8 master mold on silicon, with the first layer having a thickness of ∼4 μm, and the second layer ∼96 μm. These master molds were then used to produce microfluidic devices using polydimethylsiloxane (PDMS, Dow Sylgard 184 Silicone elastomer base and curing agent). These devices were affixed to 30mm apertured glass bottom dishes (WillCo Wells, GWST-5030). Ngn2 Lower induced motor neurons were dissociated at day 5 and plated in compartmentalized microfluidic devices mounted on glass-bottom dishes coated with 0.1 mg/ml poly-L-ornithine (Sigma-Aldrich) diluted in 50 mM Borate buffer, pH=8.5 and 5 µg/ml laminin (Invitrogen) at a concentration of around 75,000 neurons/device.

#### Axotomy

Axonal injury was performed at day 14 and day 42 of culture by repeated vacuum aspiration and reperfusion with 1X PBS in the axon chamber until axons were cut effectively without disturbing cell bodies in the soma compartment. To visualize repaired axons, devices were fixed at 24-, 48-, and 72-hours post-injury and stained for βIII tubulin following the previously described immunostaining protocol. 2 independent replicates were performed for DIV14 and DIV42 experiments, the range of axons measured for the whole experiment per genotype were: wild-type (103-249 axons at DIV14, 223-283 axons at DIV42) and *KIF5A^null^* (24-181 axons at DIV14, 46-53 at DIV42). For measuring the length or recovered axons, axons were traced from site of injury (end of microchannels) and length was measured with the NeuronJ^116^ plugin from Image (NIH).

### Live Cell Imaging and trafficking analysis

Live imaging experiments were performed on a Nikon Ti-E Eclipse inverted microscope system fitted with a Confocal A1R (LU4A Laser) unit and an EMCCD camera (Andor DU-897). The inverted microscope together with the EMCCD were used for live imaging using the LU4A laser unit. Neurons in microfluidic devices were placed in a Tokai Hit stage-top incubator (STRG-WELSX-Set, Tokai Hit, Japan) and imaged at 37C, 5% CO2 with humidity through the bath unit heater in neuronal maintenance medium (Life Technologies # 11570556) including 200mM Glutamax, 20% Dextrose, Non-Essential Amino Acids (NEAA; Life Technologies # 11350912), B27 (Life Technologies # 17504044, 0.1% P/S, 0.5% Fetal bovine serum (Life Technologies # 10270106), 10ng/ml BDNF (Peprotech # 450-10), 10ng/ml CNTF (Stem Cell Technologies # 78010.1) and 10ng/ml GDNF (Peprotech # 450-10). Mitochondria and neurofilaments were imaged using a 40x oil objective lens (NA 1.3, Nikon) while Halo-SFPQ granules were imaged using a 60x oil objective lens (NA 1.4, Nikon), with a Perfect Focus System. Time lapse recordings were acquired at a framerate of 1 second per frame for 5 minutes for MitoTracker-CMX Ros (#M7512), 0.5 second per frame for 5 minutes for Halo-SFPQ granules and 3 seconds for 10 minutes for Neurofilaments^105^ (NEFL-GFP,. For neurofilament transport, thinner axons were imaged and regions of 180µm length were photobleached based on guidelines established by Anthony Brown’s lab^58^ (, that allow for detection of moving polymers without photodamage and enable long moving filaments to be recorded. Thin axons were photobleached for 1 minute before acquisition (laser intensity and pulse duration were optimized to reach >90% fluorescence decrease of GFP) and images were acquired for 660 seconds after an initial 5 seconds of baseline.

Kymographs were generated using the KymographBuilder plugin for Fiji (line width of 25-50, spanning the width of the microchannels). Tracks of individual organelles/filaments/granules were manually traced using Fiji. Lines fitted over tracks in the kymographs were analyzed with a custom-made Kymograph Analyzer tool ImageJ macro (available upon request). The macro extracts the dynamic parameters directly from the individual trajectories (change in space and time, speed). These parameters are loaded in and analyzed with a custom made MATLAB script which is publicly available (https://github.com/alemoro/FGA_LCI_tools/blob/masters/KymoData_scripts.m). Motile organelles/filaments/granules (anterograde or retrograde) were defined by net displacement >2 μm while nonmotile (stationary) ones were defined by net displacement <2 μm. The number values (tracks) reported in the supplemental figures 3 and 4 represent the number of segments (motile and non-motile) analyzed within each replicate for each cargo, genotype and time.

### Quantification and Statistical Analysis

Graphs were generated using GraphPad Prism (v. 10). In figures, boxplots show the median value, interquartile range, and whiskers including all values within 1.5 times IQR from the median (Tukey-style whiskers) or whiskers from minimum to maximum values. No outliers were removed. In other figures, bars and lines represent the mean and standard deviation. Equal variances were tested using Brown-Forsythe test with a *p* value < 0.05 as considered significant. Statistical analyses were performed using a two-tail unpaired Student’s t-test, One-Way or Two-Way ANOVAs with a *p* value of < 0.05 considered as significant using GraphPad Prism 10.0. Post-HOC analysis following significant ANOVAs were determined based on whether the data was parametric. Normally distributed data utilized a Tukey post-HOC test, while non-parametric data utilized Kruskal-Wallis’ multiple comparison test for One-Way ANOVA. Tests between different genes across multiple age groups or two time points used Two-Way ANOVA with Sidak’s multiple comparison test.

## Supplemental materials

Supplemental table 1: Differential expression analysis

Supplemental table 2: Significantly regulated proteins in PantherDB protein families

Supplemental table 3: Detected proteins

Supplemental file 1: Photomask design

Supplemental file 2: Time-lapse images of trafficking data represented in figures.

## Acknowledgements

Support to K.E. was provided by Target ALS, NIH 5R01NS089742 and the Harvard Stem Cell Institute. I.G.S.J. is supported by Target ALS. The authors thank Franks Koopmans for assistance and help with analysis of proteomics data and the Department of Molecular and Cellular Neurobiology for infrastructure and support with MS experiments. We thank Desiree Schut, Judith Huijgen, Solange Lopes Cardozo and Jennifer Vlaardingerbroek for assistance in hESC culture and maintenance of hESC-derived neurons and glia cultures; Robbert Zalm and Ingrid Saarloos for cloning and production of lentiviral particles as well as insight and expertise in protein chemistry experiments. We thank Dean De Boer for design and production of microfluidic devices. We thank Alessandro Moro for developing scripts to analyze the transport data and Rein Hoogstraaten for support on photobleaching experiments. We thank Francesco Limone for insight and expertise on the Ngn2 motor neuron differentiation pipeline. We thank Amparo Roig for assistance in analysis of BrainSpan datasets. Schematic diagrams in all Figures were created with BioRender.com.

## Authors Contributions

Conceptualization, I.G.S.J., M.V., R.F.T., K.E.; Methodology, I.G.S.J., J.B.; Validation, I.G.S.J., J.B.; Investigation, I.G.S.J., J.B.; Formal analysis, I.G.S.J., J.B.; Data Curation, I.G.S.J., J.B., M.V., R.F.T; Resources, M.V., R.F.T, K.E.; Writing - Original Draft, I.G.S.J.; Writing - Review & Editing, I.G.S.J., M.V., R.F.T; Visualization I.G.S.J., J.B.; Supervision, I.G.S.J., M.V., R.F.T; Project Administration, M.V., R.F.T, KE; Funding Acquisition, M.V., K.E.

